# Centromeric localization of KNL2 and CENP-C proteins in plants depends on their centromere-targeting domain and DNA-binding regions

**DOI:** 10.1101/2024.04.11.588992

**Authors:** Surya Prakash Yalagapati, Ulkar Ahmadli, Aditya Sinha, Manikandan Kalidass, Siarhei Dabravolski, Sheng Zuo, Ramakrishna Yadala, Twan Rutten, Alexandre Berr, Paul Talbert, Inna Lermontova

**Affiliations:** Leibniz Institute of Plant Genetics and Crop Plant Research (IPK) Gatersleben, Corrensstrasse 3, D-06466 Seeland, Germany; Department of Biotechnology Engineering, Braude Academic College of Engineering, Snunit 51, Karmiel 2161002, Israel; Anhui Provincial Key Laboratory of the Conservation and Exploitation of Biological Resources, College of Life Sciences, Anhui Normal University, Wuhu, 241000, China; Institut de Biologie Moléculaire des Plantes (IBMP), CNRS, Université de Strasbourg, Strasbourg, France; Howard Hughes Medical Institute, Basic Sciences Division, Fred Hutchinson Cancer Research Center, Seattle, WA 98109, USA

**Keywords:** Centromere targeting, KNL2, CENPC, CENPC-k motifs, CENPC motifs, DNA-binding regions

## Abstract

In eukaryotic organisms, proper chromosome segregation during cell division depends on the centromeric histone H3 (CENH3) variant. Our previous studies identified a plant CENH3 assembly factor, Kinetochore Null2 (αKNL2), that possesses a centromere-targeting motif, CENPC-k, similar to the CENPC motif in CENP-C. Additionally, we have demonstrated that αKNL2 can bind DNA *in vitro,* independent of its CENPC-k motif. Thus, the mechanism underlying the binding of αKNL2 to centromeric DNA remains elusive.

Our study shows that the CENPC-k and CENPC motifs alone are not sufficient to target the centromere in *N. benthamiana* and *A. thaliana*. *In-silico* analysis revealed flanking DNA-binding regions near the CENPC-k and CENPC motifs, suggesting their importance in interacting with centromeric DNA. Fusion of protein fragments containing these motifs to EYFP facilitated targeting to the centromere. Deletion of DNA-binding domains reduced the centromeric localization of αKNL2-C, whereas fusion of CENPC-k to the H-NS protein from E. coli targeted it to centromeres.

We conclude that targeting of αKNL2 and CENP-C proteins to centromeres is dependent on the CENPC-k/CENPC motifs, and their sequence-independent DNA-binding promotes anchoring at the centromere. Understanding the targeting mechanisms of KNL2 and CENP-C may help to engineer kinetochore structure by targeting chromatin modifying proteins to centromeres.

## Introduction

Centromeres are specific chromosomal positions where the kinetochore complex is established for the proper attachment of the spindle microtubules and the correct separation of chromosomes during mitosis and meiosis. Centromeric DNA sequences are highly variable even among closely related species and are composed of centromeric repeats and transposable elements (Lermontova *et al*., 2014, Lermontova *et al*., 2015). These sequences are neither required nor sufficient for centromere formation as so-called neocentromeres can be formed at atypical chromosomal regions such as chromosome arms or telomeres (reviewed by Scott and Sullivan (2014)). In centromeric chromatin, a significant proportion of nucleosomes is characterized by the presence of the centromere-specific histone H3 variant CENH3 instead of canonical H3. The presence of CENH3 homologs in all animals, fungi, and plants studied so far has resulted in an assumption that CENH3-containing chromatin is a basic requirement for centromere function (Malik and Henikoff, 2003, Panchenko and Black, 2009). However, Drinnenberg *et al*. (2014) have demonstrated that several lineages of holocentric insects contain a CENH3-independent centromere. Loading of CENH3 into centromeric nucleosomes is a tightly regulated process that can be divided into three steps: licensing of the centromere, loading of CENH3 by histone chaperones, and stabilization of the newly incorporated CENH3 in centromeric nucleosomes (Lermontova *et al*., 2013). The role of centromere-licensing factors was proposed for the Mis18 complex containing Mis18α, Mis18β, M18BP1/Kinetochore Null2 (KNL2) proteins (Fujita *et al*., 2007). Of this complex, only Mis18 proteins (Mis16, Mis19 (Eic1), and Mis20 (Eic2) was found in yeast (Hayashi *et al*., 2004), and only M18BP1/KNL2 - in *C. elegans* (Maddox *et al*., 2007) and plants (Lermontova *et al*., 2013). Reduced *KNL2* expression in plants results in reduced level of CENH3 at centromeres of meristematic nuclei, anaphase bridges during mitosis, micronuclei in pollen tetrads followed by reduced growth rate and fertility (Lermontova *et al*., 2013). Recently it was shown that in plants the *KNL2* gene underwent three independent ancient duplications, namely in ferns, grasses, and eudicots (Zuo *et al*., 2022). Additionally, previously unclassified *KNL2* genes could be divided into the clades αKNL2 and βKNL2 in eudicots and γKNL2 and δKNL2 in grasses, respectively. Thus, in this study, *Arabidopsis* KNL2, which was previously reported, will be referred to as αKNL2 (Lermontova *et al*., 2013). All KNL2 homologs identified so far are characterized by the conserved N-terminal SANTA (SANT associated) domain. In vertebrates, the SANTA domain is present in parallel with the putative DNA-binding domain SANT. In contrast, in both KNL2 variants of plants, the SANT domain cannot be detected (Lermontova *et al*., 2013, Zuo *et al*., 2022), although the C-terminal part of αKNL2 demonstrated DNA-binding capacity (Sandmann *et al*., 2017).

A conserved CENPC-k motif identified in α- and γKNL2 of plants and in KNL2 of non-mammalian vertebrates (Kral, 2016, Sandmann *et al*., 2017) was not found in β- and δKNL2 variants of plants and in mammals (Zuo *et al*., 2022). The CENPC motif was originally considered as a typical feature of the CENP-C protein required for its targeting to the centromere and interaction with the CENH3 nucleosome (Kato *et al*., 2013). Later, the CENPC-k motif was shown to be similarly required for targeting KNL2 protein homologs to centromeres in *Arabidopsis* (Sandmann *et al*., 2017), chicken (Hori *et al*., 2017), and frog (French *et al*., 2017). In addition, a recent study by Jiang *et al*. (2023) showed that the CENPC-k motif of chicken KNL2 is able to bind directly to the CENH3 in centromeric nucleosome. For new CENH3 deposition in human cells, M18BP1/KNL2, which lacks the CENPC-k motif (Kral, 2016), localizes to centromeres via CENP-C binding (McKinley and Cheeseman, 2014, Nardi *et al*., 2016). In contrast to this in *Xenopus* egg extract or chicken DT40 cells, association of KNL2 containing the CENPC-k motif (Kral, 2016) with interphase centromeres does not require CENP-C (Perpelescu *et al*., 2015, Westhorpe *et al*., 2015). Despite the low sequence similarity between KNL2 and CENP-C, the two proteins share the presence of a CENPC-like motif and the ability to bind DNA and RNA molecules. Both αKNL2 and CENP-C proteins of plants can bind DNA in a sequence-independent manner *in vitro* while *in vivo,* they are preferentially associated with the centromeric repeats (Du *et al*., 2010, Sandmann *et al*., 2017). Additionally, human CENP-C have shown interaction with DNA in a similar way (Politi *et al*., 2002). The fact that the C-terminal fragment of αKNL2 can target centromeres of *N. benthamiana*, even though its centromeric sequences differ from those of *A. thaliana*, further confirms that the targeting of KNL2 to centromeres and its binding to DNA does not depend on the sequence of centromeric DNA (Sandmann *et al*., 2017).

Here we show that the expression of fusion constructs of CENPC-k and CENPC motifs of αKNL2 and CENP-C proteins, respectively, with EYFP resulted in a distribution of fluorescence in the nucleoplasm and cytoplasm of transiently transformed *N. benthamiana* and stably transformed *A. thaliana*, demonstrating that these motifs alone are insufficient for targeting centromeres. We identified putative DNA-binding regions near the CENPC motifs of αKNL2 and CENP-C proteins and demonstrated that protein fragments containing CENPC motifs and DNA-binding sites can target centromeres. Moreover, deleting one of the DNA-binding sites from the C-terminus of αKNL2 showed a reduced localization to centromeres, whereas the deletion of both abolished its centromeric localization. Using *in-silico* analysis and EMSA assay, we have also demonstrated that the deletion of putative DNA-binding regions reduces the interaction of αKNL2-C with the centromeric repeat *pAL1*. Targeting of protein fragments containing the CENPC-k motif in combination with bacterial DNA-binding region(s) to centromeres showed that any DNA-binding regions with sequence-independent DNA-binding ability are sufficient to anchor kinetochore proteins to centromeres.

## Materials and Methods

### Plasmid construction and site-directed mutagenesis

All fragments including the CENPC-k and CENPC motifs with and without DNA-binding regions (Supplementary Table 1-4) were amplified from the αKNL2 and CENP-C cDNA clones in pDONR221 (Invitrogen) vector using primers listed in Supplementary Table 5. Subsequently, all fragments were separately cloned into the pDONR221 vector via the Gateway Cloning (BP reaction). To delete the upstream DNA-binding region (468-477) of the CENPC-k motif of αKNL2, the αKNL2-C/pDONR221 construct (Lermontova et al., 2013) was subjected to PCR mutagenesis using the Phusion site-directed mutagenesis kit (Thermo Fisher Scientific) and the following primer pairs αKNL2-CΔDNAb (1)f and αKNL2-CΔDNAb (1)r were used. To delete the downstream DNA-binding region (581-598) of the CENPC-k motif, the αKNL2-C fragment was amplified from the αKNL2-C in pGWB41 vector using αKNL2-CΔDNAb (2)f and αKNL2-CΔDNAb (2)r primers listed in Supplementary Table 5 and cloned subsequently into the pDONR221 vector. All plasmid clones were prepared for sequence analysis using the QIAGEN QIAprep kit according to the instructions of the supplier (QIAGEN). From pDONR221 clones, the open reading frames were recombined via Gateway LR reaction (Invitrogen) into the two attR recombination sites of the Gateway-compatible vectors pGWB641, pGWB642 and pGWB441 (https://shimane-u.org/nakagawa/), respectively, to study the localization of proteins *in-vivo*. To assess the influence of a non-plant DNA binding domain on the centromeric localization of the CENPC-k motif, the non-specific DNA binding domain (Nsdbd) was isolated from the bacterial histone-like nucleoid structuring protein (H-NS). Firstly, the genomic DNA from *Escherichia coli* was purified following the Wizard Genomic DNA Purification Kit (Promega) protocol and served as a template for PCR amplification. The Nsdbd was then amplified from this template using primers listed in Supplementary Table 5. Following gel purification, the PCR product was ligated into the pGEM-T Easy vector (Promega). Subsequently, after validation through sequencing, the Nsdbd fusion was re-amplified with Gateway-compatible primers (Supplementary Table 5) and cloned into the pDONR221 vector (Invitrogen). Finally, the Nsdbd fragment was recombined from a pDONR221 clone through Gateway LR reaction into the Gateway-compatible vector pB7WGF2 (Karimi *et al*., 2002). To generate fusion constructs of CENPC-k motif with Nsdbd DNA fragments of ATG-Nsdbd-CENPC-k-Nsdbd and ATG-Nsdbd-CENPC-k (Supplementary Table 4) were synthesized by BioCat company (https://www.biocat.com/) and provided as clones in pDONR221 vectors. From pDONR221 clones, the open reading frames were recombined via Gateway LR reaction (Invitrogen) into the two attR recombination sites of the Gateway-compatible vectors pGWB641, pGWB642.

### Plant transformation and cultivation

*Agrobacterium* tumefaciens-mediated transient transformation of *Nicotiana benthamiana* plants was performed according to (Walter *et al*., 2004). After two days of infiltration, epidermal cell layers of tobacco leaves were assayed for fluorescence and at least 50 cells per plant were analyzed. For each construct, transient transformation of *N. benthamiana* was repeated at least three times.

Plants of *Arabidopsis* accession Columbia-0 were transformed according to the floral dip method (Clough and Bent, 1998). T1 transformants were selected on Murashige and Skoog (MS) medium (Murashige and Skoog, 1962) containing 20 mg/L of phosphinotricine (transformants with pGWB641, pGWB642 vectors) or 40 mg/L of kanamycin (transformants with pGWB441 and pGWB442 vectors). Growth conditions in a cultivation room were 21°C, 8 h light/18°C, 16 h dark or 21°C, 16 h light/18°C, 8 h dark.

### Microscopy analysis of fluorescent signals

Immunostaining of nuclei/chromosomes was performed according to (Ahmadli *et al*., 2022). For live cell imaging, *Arabidopsis* seeds harboring fusion constructs were germinated on agar medium. Roots of 7-day-old seedlings were investigated with a LSM780 confocal laser microscope (Carl Zeiss, Jena, Germany) using a 40x/1.2 water immersion objective. EYFP was excited with a 488-nm laser line and fluorescence analyzed with a 505-550 nm band-pass. Images were recorded with the LSM software zen black release 3.2.

### In-silico prediction of the DNA-binding ability of proteins

To predict the DNA-binding ability of M18BP1/KNL2 proteins of plants, mammalian and non-mammalian vertebrates, and C.elegans, structural models were constructed using the SWISS-MODEL (Waterhouse *et al*., 2018) for Q7XVZ3, <u>B8A9Q3, A0A8M2B5N0, A0A8V0Z4U5</u> or down-loaded from AlphaFold Protein Structure Database (https://alphafold.ebi.ac.uk/) (Jumper *et al*., 2021, Varadi *et al*., 2022) for other proteins. The presence of DNA-binding regions withing selected proteins was detected with the following sequence- and structure-based on-line tools: DNABIND (Liu and Hu, 2013), DRNApred (Yan and Kurgan, 2017), DNAgenie (Zhang *et al*., 2021), and NucBind (Su *et al*., 2019).

### Homology modelling and protein-DNA interaction by molecular docking

The protein sequence of the centromeric protein αKNL2 was retrieved from the UniProtKB database (https://www.uniprot.org/) in FASTA format. Homology modeling of the αKNL2-C and CENP-C proteins using the FASTA sequence was performed using I-TASSER (Iterative Threading ASSEmbly Refinement). I-TASSER is a hierarchical approach to predicting protein structure and structure-based function annotation. The program detects structural templates from the PDB (Protein Data Bank) using a multiple threading method (LOMETS), then builds full-length atomic models using iterative template-based fragment assembly simulations. I-TASSER is available at https://zhanggroup.org/I-TASSER/.

HDOCK is an online server used to identify the protein-protein and protein-DNA/RNA docking based on a hybrid algorithm of template-based modeling and *ab initio* free docking (http://hdock.phys.hust.edu.cn/). For the centromeric DNA sequence, the *pAL1* repeat sequence has been retrieved. Modeled PDB structures were submitted to the HDOCK server for the protein-DNA interaction. The interaction between two molecules is determined based on the two scores, as docking score and the confidence score. A more negative docking score means a possible binding model. In accordance with the confidence score, the two molecules would be very likely to bind if the score is above 0.7, the two molecules would be possible to bind when the score is between 0.5-0.7, and if it is below 0.5, the two molecules would be unlikely to bind.

### Electrophoretic mobility shift assay

The pF3A WG (Promega) vectors containing the CDS of αKNL2-C, αKNL2-CΔDNAb (1), αKNL2-CΔDNAb (2) and αKNL2-CΔDNAb (1,2) fused with a FLAG tag were used for protein expression with TNT SP6 High Yield Wheat Germ (Promega) following the manufacture’s protocol, incubating the reaction for 2 h at 25°C. Expression of proteins was validated by performing a Western blot analysis against the FLAG-epitope tag using an anti-FLAG antibody (Sigma). For the EMSA assays, the DNA fragment (*pAL1*) was amplified by PCR using fluorocently labelled (IR-DYE-700) primers, and the probe was purified using oligonucleotide purification kit from BioRad. Binding reaction was setup using the Odyssey EMSA kit (LICOR). The wheat germ expressed proteins (3 µl) and 1 ng of probe were incubated for 30 min at RT. After incubation 2 µl of orange loading dye was added to the reaction and samples were loaded and separated on 5% native polyacrylamide gel at 4°C with 70 V till the dye reaches the bottom of the gel. Gels were imaged using LICOR Odyssey scanner.

## Results

### CENPC-k and CENPC motifs are necessary but insufficient for independent centromere targeting

The CENPC-k motif is essential for targeting the αKNL2 protein to centromeres in *Arabidopsis.* The mutagenesis of conserved amino acids or complete deletion of the CENPC-k motif abolished the centromeric localization of αKNL2. However, this the centromeric localization can be restored by replacing the CENPC-k motif of αKNL2 with the motif from the CENP-C protein (Sandmann et al., 2017). The CENPC-k motif, spanning amino acids 538-572, is present at the C-terminal part of the αKNL2 protein (Figure 1A). To test whether a CENPC-k motif alone is sufficient for centromeric targeting, we fused the CENPC-k motif to an enhanced yellow fluorescent protein (EYFP) driven by the 35S promoter. Since the fusion direction can strongly influence the subcellular localization of the protein, constructs with N- or C-terminal fusion of EYFP with the CENPC-k motif were designed (Figure 1B). To analyze the subcellular localization of EYFP-tagged CENPC-k motifs in plants, young *Nicotiana benthamiana* plants were infiltrated with *Agrobacterium* suspensions carrying the respective constructs. Two days after infiltration, fluorescent signals were detected in the nucleoplasm and cytoplasm in all cases. The fluorescence was distributed homogeneously in the nucleoplasm, and no signals corresponding to centromeres were detected (Figure 1C, D, upper panels). Subsequently, these constructs were stably transformed in *Arabidopsis*, and T1 transgenic plants were obtained. Analysis of at least ten independent transgenic lines expressing each construct showed cytoplasmic and nucleoplasmic fluorescent signals in root tips, consistent with our observations in *N. benthamiana* (Figure 1C, D, lower panels).

**Figure 1.**
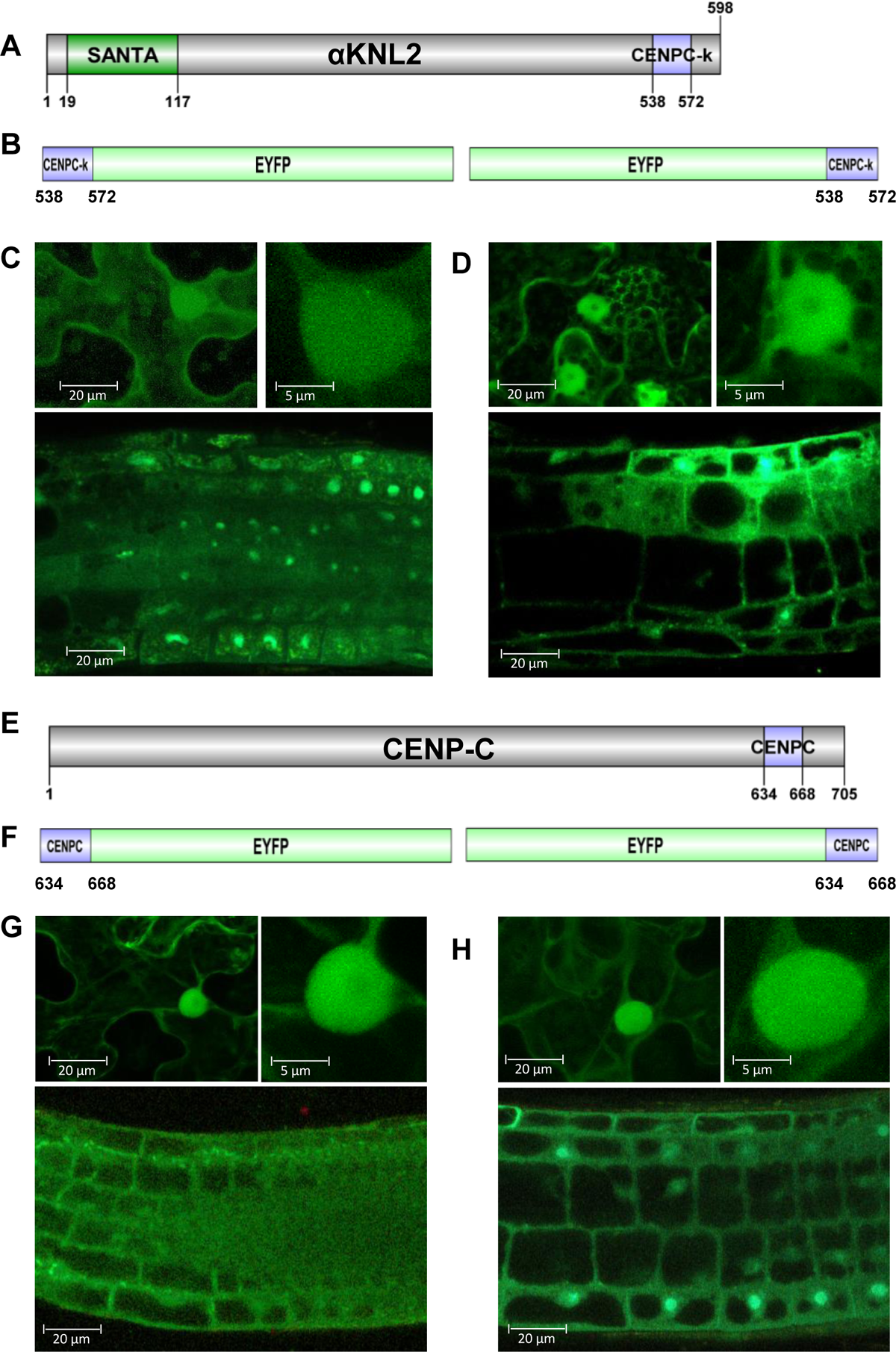
The CENPC-k and CENPC-m motifs of the KNL2 and CENP-C proteins were unable to target centromeres independently in *N. benthamiana* and *A. thaliana*. **(A)** Schematic representation of the KNL2 protein with the conserved SANTA domain and CENPC-k motif. **(B)** CENPC-k-EYFP and EYFP-CENPC-k fusion constructs **(C, D)** subcellular localization of the corresponding fusion proteins in leaves of transiently transformed *N. benthamiana* (upper parts) and roots of stably transformed *A. thaliana* (lower parts). In all cases, fluorescent signals were detected in the nucleoplasm and cytoplasm. No signals corresponding to centromeres were observed.**(E)** Schematic representation of the CENP-C protein with the conserved CENPC motif. **(F)** CENPC-EYFP and EYFP-CENPC fusion constructs. **(G, H)** The subcellular localization of the corresponding fusion proteins in transiently transformed *N. benthamiana* (upper parts) and stably transformed *A. thaliana* (lower parts). In all cases, fluorescent signals were detected in the nucleoplasm and cytoplasm. No signals corresponding to centromeres were observed.

To determine the efficiency of the CENPC motif (634-668 aa) of CENP-C in centromere targeting, this motif was fused with EYFP in both orientations (Figure 1E, F). Similar to the CENPC-k motif, the localization analysis in *N. benthamiana* revealed that both fusion variants CENPC-EYFP/EYFP-CENPC localized to both nucleoplasm and cytoplasm (Figure 1G, H, upper panels). Moreover, stable *Arabidopsis* lines harboring these constructs failed to exhibit specific signals at centromeres, and localized to nucleoplasm/cytoplasm in root tips (Figure 1G, H, lower panels).

### *In-silico* identification of DNA-binding regions in αKNL2 and CENP-C proteins

The results presented above clearly demonstrated that the CENPC-k and CENPC motifs alone are insufficient for targeting centromeres. Sandmann *et al*. (2017) showed earlier that the C-terminal region of αKNL2, harbouring the centromere-targeting CENPC-k motif, binds to centromeric DNA *in vitro* and preferentially associates with the centromeric DNA *in vivo*. Similarly, the CENP-C protein of maize has an ability to bind DNA (Du *et al*., 2010). We thus hypothesized that adding putative DNA-binding region(s) to the CENPC-k or CENPC-m motif might lead to efficient centromere targeting.

To predict putative DNA-binding regions within *Arabidopsis* αKNL2 and CENP-C proteins (Supplementary dataset 1), we employed a combination of sequence-based and structure-based predictors, including DP-Bind (Hwang *et al*., 2007), DRNApred (Yan and Kurgan, 2017), DNAgenie (Zhang *et al*., 2021) and GraphBind (Xia *et al*., 2021) programs. All predicted data consistently identified DNA-binding sites within αKNL2 and CENP-C proteins (Figure 2A, C), corroborating our previous findings. Specifically, DNA-binding regions within αKNL2 were identified at amino acid positions 468-477 (αKNL2-C-DNAb (1)), 581-598 (αKNL2-C-DNAb (2)) (Figure 2A) and in CENP-C at positions 606-617 (CENPC-DNAb (1)), 671-688 (CENPC-DNAb (2)) (Figure 2C), thus closely flanking CENPC-k and CENPC motifs, respectively. In addition, putative DNA-binding sites were also identified within the conserved CENPC-k and CENPC motifs themselves. Since deletion of the CENPC-k motif from the C-terminal part of αKNL2 did not influence its ability to interact with DNA *in vitro* (Sandmann *et al*., 2017), these DNA-binding sites may not be essential, at least for αKNL2. To have a broad view of the conservation and variation of DNA-binding regions in αKNL2 and CENPC, we produced an alignment of αKNL2 and CENPC proteins across plant species (Figure 2B, D). While numerous residues within these regions exhibited variability among homologs, the presence of conserved positively charged amino acid residues, responsible for DNA binding, was observed across nearly all homologs. This implies that despite variation, αKNL2 and CENPC retain their ability to bind DNA in plants. To test whether an ability to bind DNA/RNA is a common feature of M18BP1/KNL2 proteins, we extended our *in-silico* prediction analysis to βKNL2 of *A. thaliana*, δKNL2 of *O. sativa* and M18BP1 proteins of *H. sapiens*, *M. musculus*, *D.rerio*, *G. gallus* and *C. elegans*. This analysis using sequence- and structure-based online tools: DNABIND (Liu and Hu, 2013), DRNApred (Yan and Kurgan, 2017), DNAgenie (Zhang *et al*., 2021), and NucBind (Su *et al*., 2019) showed that all selected proteins have the ability to bind nucleic acids (Table 1).

**Figure 2.**
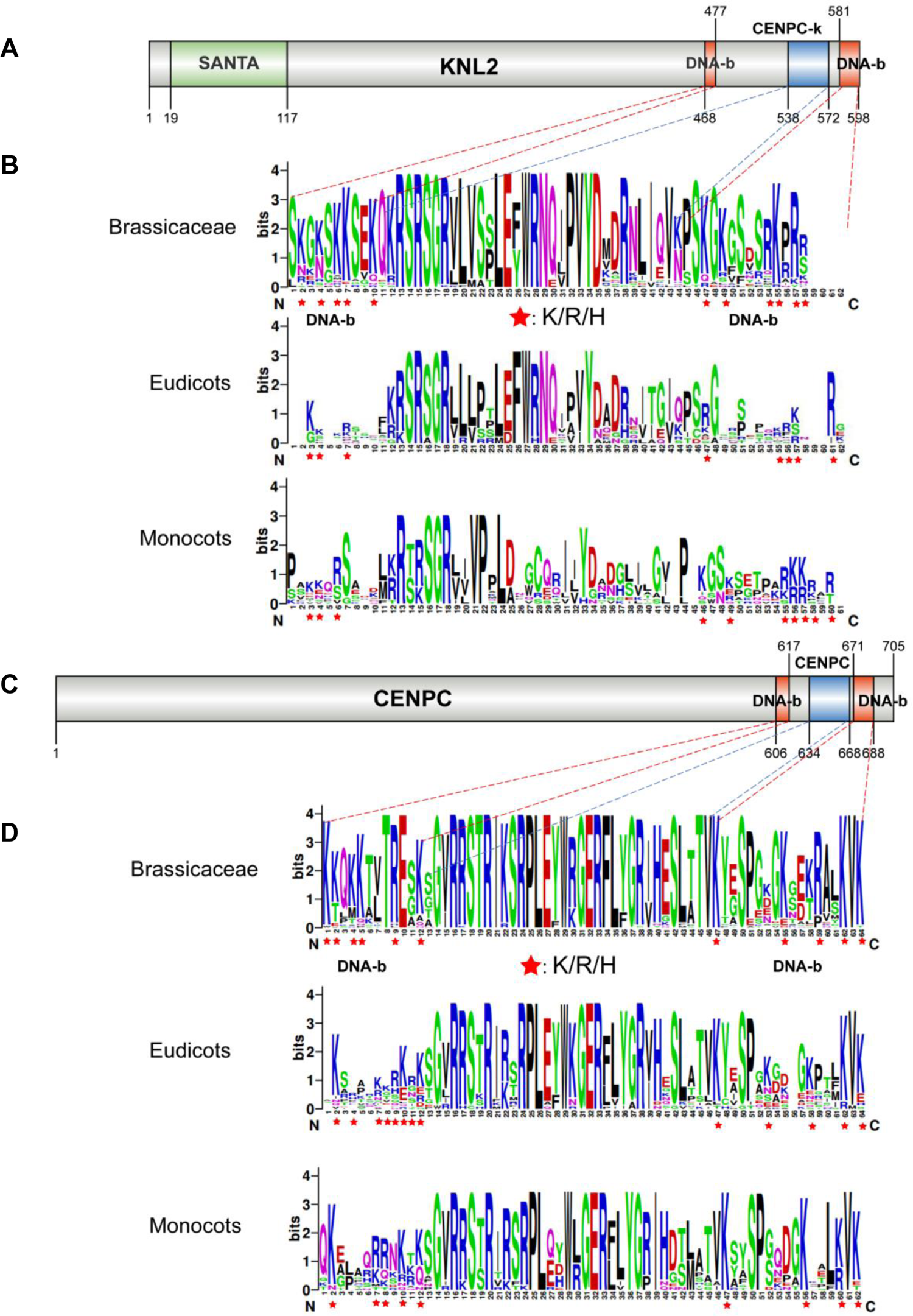
The DNA-binding motifs of KNL2 and CENP-C proteins are much less conserved than the CENPC-k motif across different plant species. **(A, C)** Domain map of KNL2 and CENP-C proteins with indicated positions of SANTA domain, CENPC-k, and DNA-binding motifs. **(B, D)** Alignment of CENPC-k, CENPC-m and DNA-binding motifs is presented in LOGO format (WebLogo; http://weblogo.berkeley.edu/logo.cgi) for easier comparison of similarities and differences in these motifs among the Brassicaceae family, Eudicot and Monocot plant species. The positively charged amino acid residues (R/K/H) of DNA-binding motifs from KNL2 and CENP-C orthologs were marked by red stars.

**Table 1.**
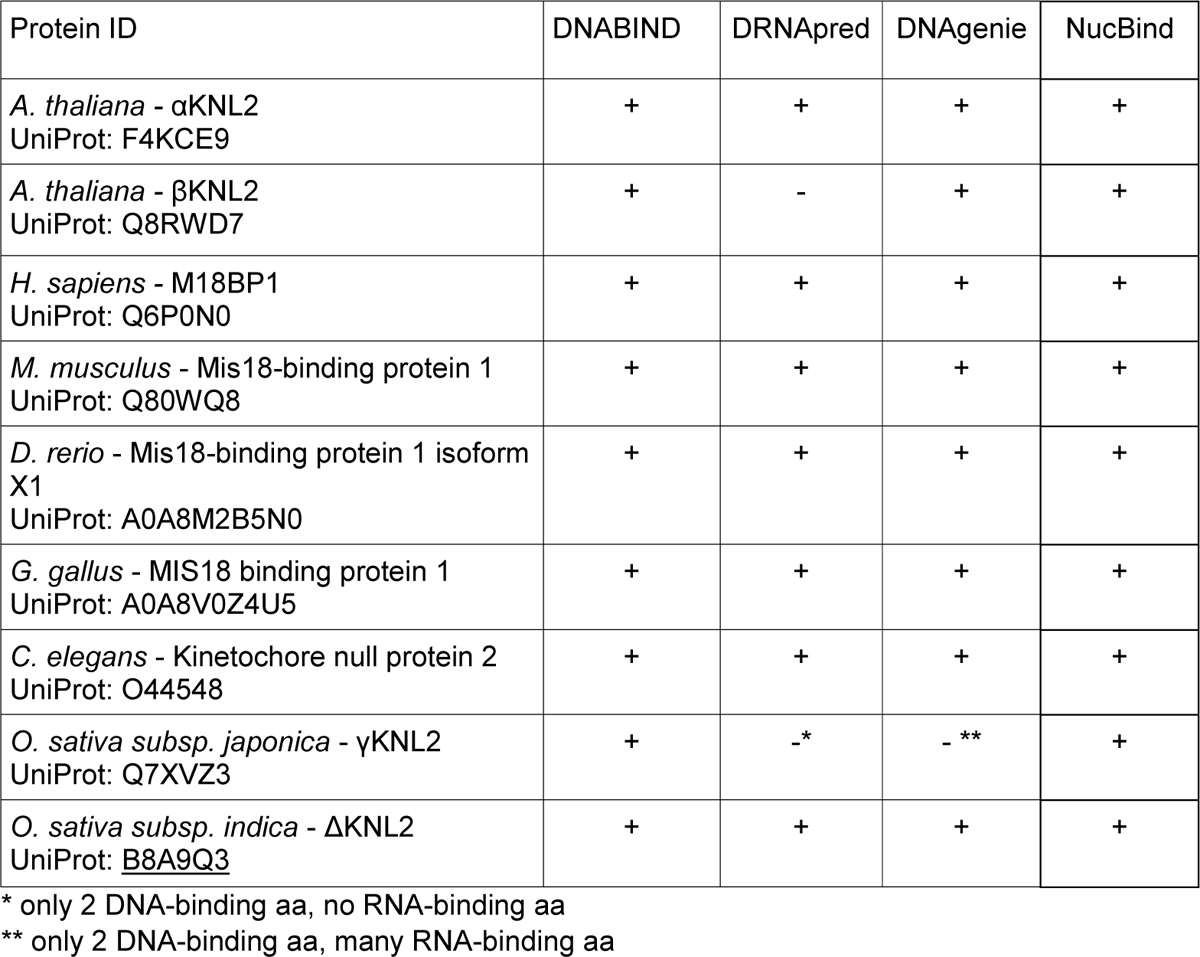
*In silico* prediction of DNA-binding regions withing KNL2 homologues.

### Molecular docking analysis predicts the interaction of αKNL2-C and CENP-C DNA-binding protein regions with centromeric DNA (*pAL1*)

Following the identification of the putative DNA binding sites on αKNL2 and CENP-C proteins, the interactions between their DNA-binding regions and centromeric DNA (*pAL1*) were investigated using molecular docking analysis. The three-dimensional structure of the αKNL2-C protein was modelled using I-TASSER, revealing a configuration of seven helices connected by four sheets via loops (Supplementary Figure 1A). To assess the interaction between αKNL2-C and *pAL1* DNA, the three-dimensional structure of αKNL2-C was docked with the *pAL1* DNA sequence using H-DOCK. This analysis revealed that the C-terminal part of αKNL2 containing the DNA-binding regions interacts with centromeric DNA. The top model showing the highest docking and confidence scores of −238.2 and 0.85, respectively, was selected for further analysis using PyMol. Our results show that both DNA-binding regions of αKNL2-C as well as a portion of the CENPC-k motif were interacting with centromeric DNA (Figure 3A). Subsequently, the αKNL2-C protein structure was modelled after the deletion of both DNA-binding regions. Upon docking αKNL2-CΔDNAb (1,2) with centromeric DNA, the docking and confidence scores were reduced to −179.1 and 0.53, respectively, with only a partial interaction observed between the CENPC-k motif and DNA (Figure 3B, Supplementary Figure 1C). Thus, the DNA-binding regions near the CENPC-k motif may play a crucial role in mediating the interaction with DNA and anchoring αKNL2 to centromeres.

**Figure 3.**
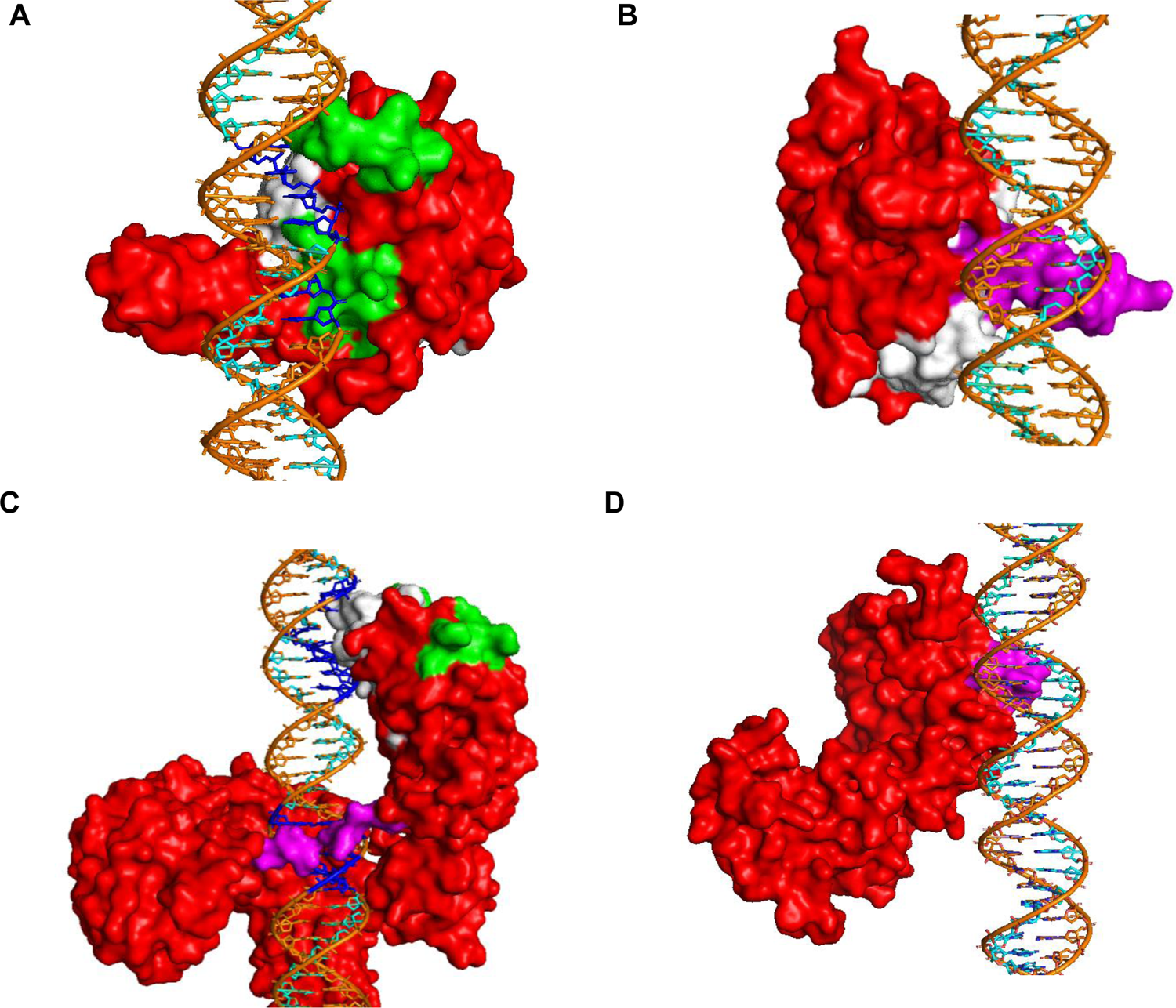
Interaction of the centromeric DNA with KNL2-C and CENPC proteins by molecular docking. **(A)** The docking of αKNL2-C (red) with centromeric DNA (*pAL1*) (orange) showing the interaction (blue) via DNA-binding regions (green) and CENPC-k motif (white). **(B)** The interaction is reduced between the αKNL2-CΔDNA-binding (1,2) (red) and *pAL1* (orange). The CENPC-k motifs (white) and some regions (magenta) interact with DNA. **(C)** The interaction of CENPC (red) with centromeric DNA (*pAL1*) (orange). The CENPC-m motif (white), DNA-binding regions (green) and some unknown regions (magenta) interact (blue) with DNA. **(D)** The reduced interaction was observed between the CENPCΔCENPCmotif, DNA-binding (red) and *pAL1* (orange). The protein is shown in surface and the DNA in cartoon sticks models.

The docking analysis was also carried out for the I-TASSER-modelled CENP-C protein with centromeric DNA (*pAL1*). The modelled CENP-C protein exhibits 22 helixes and ten sheets interconnected by loops (Supplementary Figure 1B). Results from the top predicted model reveal an interaction between the CENPC protein and centromeric DNA, with docking and confidence scores of −248.4 and 0.87, respectively. Additionally, we observed that the CENPC motif directly interacts with centromeric DNA, while DNA-binding regions may facilitate the binding of the CENPC motif to DNA (Figure 3C). Subsequently, upon deletion of both, the CENPC motif and DNA-binding regions, the docking and confidence scores were decreased to −171.8 and 0.60, respectively (Figure 3D, Supplementary Figure 1D). Thus, it is likely that the CENPC motif and the adjacent DNA-binding regions play a crucial role in the interaction with centromeric DNA.

### CENPC-k and CENPC motifs together with their DNA-binding regions can target centromeres

The bioinformatics analysis revealed that the DNA-binding regions of αKNL2 and CENP-C proteins near the CENPC-k and CENPC motifs, respectively, may be required for their interaction with centromeric DNA. To test whether the addition of DNA-binding regions to the CENPC-k or CENPC motif would result in efficient targeting to the centromere, fusion constructs containing either a αKNL2 fragment with the CENPC-k motif and putative DNA-binding regions (CENPC-k-DNAb, Figure 4A) or a CENP-C fragment with the CENPC motif and putative DNA-binding regions (CENPC-DNAb, Figure 4C) with EYFP in either direction were constructed.

**Figure 4.**
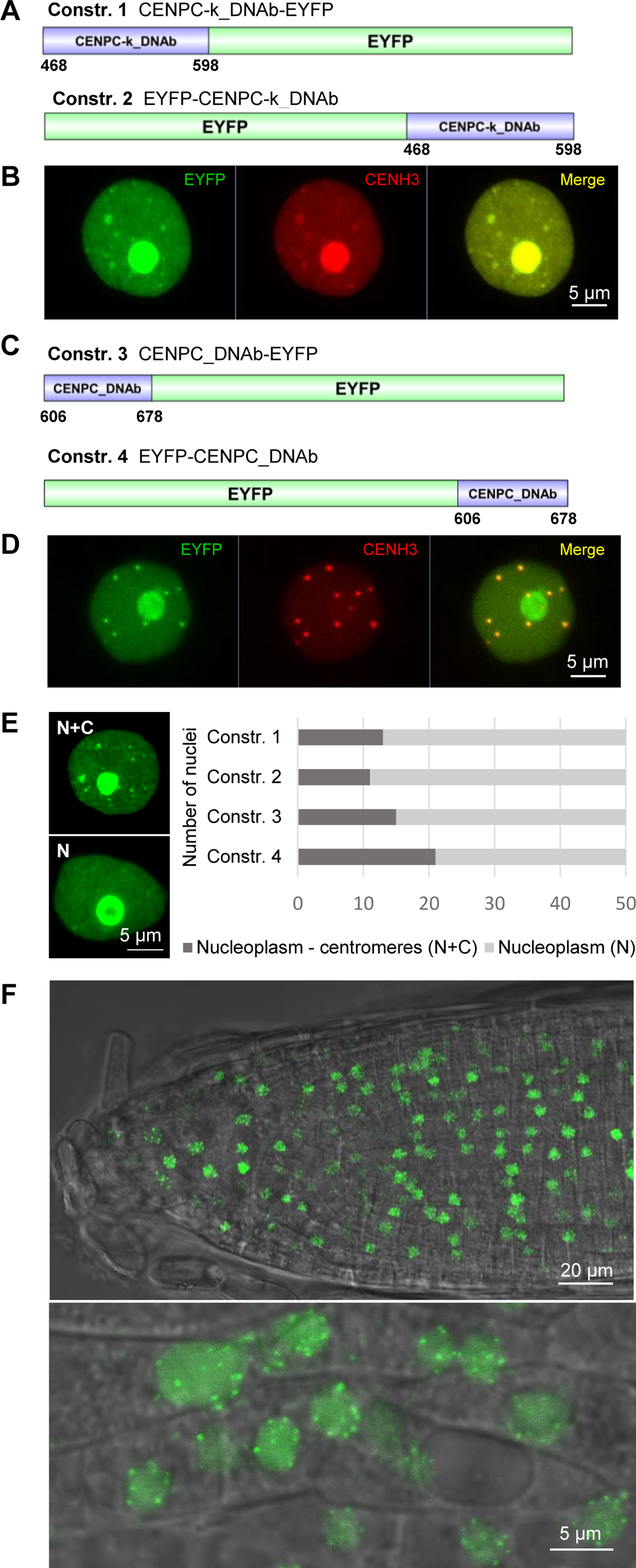
KNL2-C and CENPC containing the CENPC-k, CENPC-m motifs and the DNA-binding regions showed centromere-targeting. **(A)** The fusion constructs of CENPC-k_DNAb-EYFP (upper) and EYFP-CENPC-k_DNAb (lower). **(B)** A representative image showing the subcellular localization of the corresponding fusion proteins in the nucleoplasm, at the presumed centromeres, and in the nucleoli of transiently transformed *N. benthamiana*. **(C)** The fusion constructs of CENPC_DNAb-EYFP (upper) and EYFP-CENPC_DNAb (lower). **(D)** A representative image showing the subcellular localization of the corresponding fusion proteins in the nucleoplasm, at the presumed centromeres, and in the nucleoli of transiently transformed *N. benthamiana*. To confirm a centromeric localization of all fusion proteins in *N. benthamiana,* a co-infiltration with CENH3-mcherry construct was performed (B and D). (**E**) Distribution of fluorescence patterns in nuclei of *N. benthamiana* leaves transformed with constructs containing CENPC-k, and CENPC with DNA-binding motifs in fusion with EYFP. The two different patterns of fluorescence signals such as nucleoplasmic and centromeric (N+C) and nucleoplasmic signals (N) were defined. Frequency of nuclei displaying these two fluorescence patterns were determined for each construct based on 50 nuclei in *N. benthamiana* leaves. **(F)** A representative image showing the subcellular localization of CENPC-k_DNAb-EYFP and EYFP-CENPC-k_DNAb fusion proteins in the nucleoplasm and at the presumed centromeres of root tip nuclei of stably transformed *A. thaliana*.

Initially, all four constructs were transiently expressed in *N. benthamiana*. The *in vivo* fluorescence localization analysis in young leaves of *N. benthamiana* showed that the addition of DNA binding regions to the CENPC-k/CENPC-m motif resulted in centromere-specific signals in addition to the fluorescence distributed in the nucleoplasm (Figure 4B, D). Interestingly, fluorescent signals were consistently detected in the nucleoli of most nuclei for all constructs. However, when expressing the CENPC-k and CENPC EYFP fusion constructs in *N. benthamiana*, fluorescence within the nucleolus was only rarely observed (Figure 1G, H). This led us to speculate that the inclusion of nucleic acid binding regions may have facilitated the interaction of resulting fragments with RNA. Indeed, αKNL2 as well as CENPC exhibited the capability to bind RNA (Du *et al*., 2010, Sandmann *et al*., 2017). To confirm the centromere-specific signals, a co-localization experiment with the centromere-marker CENH3 fused to mCherry was performed. As a result, the fluorescence of CENH3-mCherry in centromeres co-localized with discrete EYFP signals of CENPC-k-DNAb/CENPC-DNAb (Figure 4B, D). Further, we analyzed the distribution of fluorescent signals in 50 nuclei of *N. benthamiana* for each fusion construct. In all cases, localization patterns in nucleoplasm and centromeres (N+C) and only in nucleoplasm (N) were observed at a similar proportion (Figure 4E).

To analyze the ability of CENPC-k-DNAb/CENPC-DNAb constructs to target the centromere in *Arabidopsis*, all four constructs were stably transformed in *Arabidopsis* plants. However, no transgenic plants were obtained with the CENPC-DNAb constructs in three repeated experiments, while the transformation performed with CENPC-k-DNAb constructs as a positive control led to the generation of transgenic lines (Supplementary Figure 2). In a case of CENPC-k-DNAb, at least ten independent transgenic lines were analyzed for each construct. In all cases, fluorescent signals were distributed in the nucleoplasm and at presumable centromeres (Figure 4F). A double immunostaining experiment with anti-GFP and anti-CENH3 antibodies on the root tip nuclei of transformants confirmed the centromeric localization of EYFP fluorescent signals (Supplementary Figure 3). Thus, the addition of DNA-binding regions to CENPС-k motif results in efficient localization to centromeres.

### Deletion of DNA-binding regions adjacent to the CENPC-k motif abolishes centromeric localization of αKNL2-C

We demonstrated that two DNA-binding regions adjacent to the CENPC-k motif can target *N. benthamiana* and *A. thaliana* centromeres. Therefore, to investigate the impact of potential DNA-binding region(s) on the centromeric localization of αKNL2, constructs encoding the C-terminal part of αKNL2 lacking one or both of the DNA-binding regions were generated and identified as αKNL2-CΔDNAb (1), αKNL2-CΔDNAb (2) and αKNL2-CΔDNAb (1,2), respectively (Figure 5A). The C-terminus was chosen instead of the full-length αKNL2 as the latter cannot be expressed in plants possibly due to its proteolytic degradation (Lermontova *et al*., 2013). All constructs were transiently expressed in *N. benthamiana*, revealing that expression of constructs with deletion of only one of the two DNA-binding regions revealed localization either in nucleoplasm and presumed centromeres, or only nucleoplasm (Figure 5B). The centromere targeting efficiency of the deletion constructs was compared with that of αKNL2-C. Analysis of 50 nuclei per sample showed that the number of nuclei with strong centromeric signals was reduced for both deletion constructs compared to the αKNL2-C. It was further observed that the decrease was more pronounced in αKNL2-CΔDNAb (2) compared to αKNL2-CΔDNAb (1), the construct with the largest deletion in the DNA-binding region. In contrast, deletion of both DNA-binding regions completely abolished the centromeric localization of the resulting protein (Figure 5C). Stable expression of these three constructs in *A. thaliana* yielded results similar to those obtained for *N. benthamiana* (Figure 5D).

**Figure 5.**
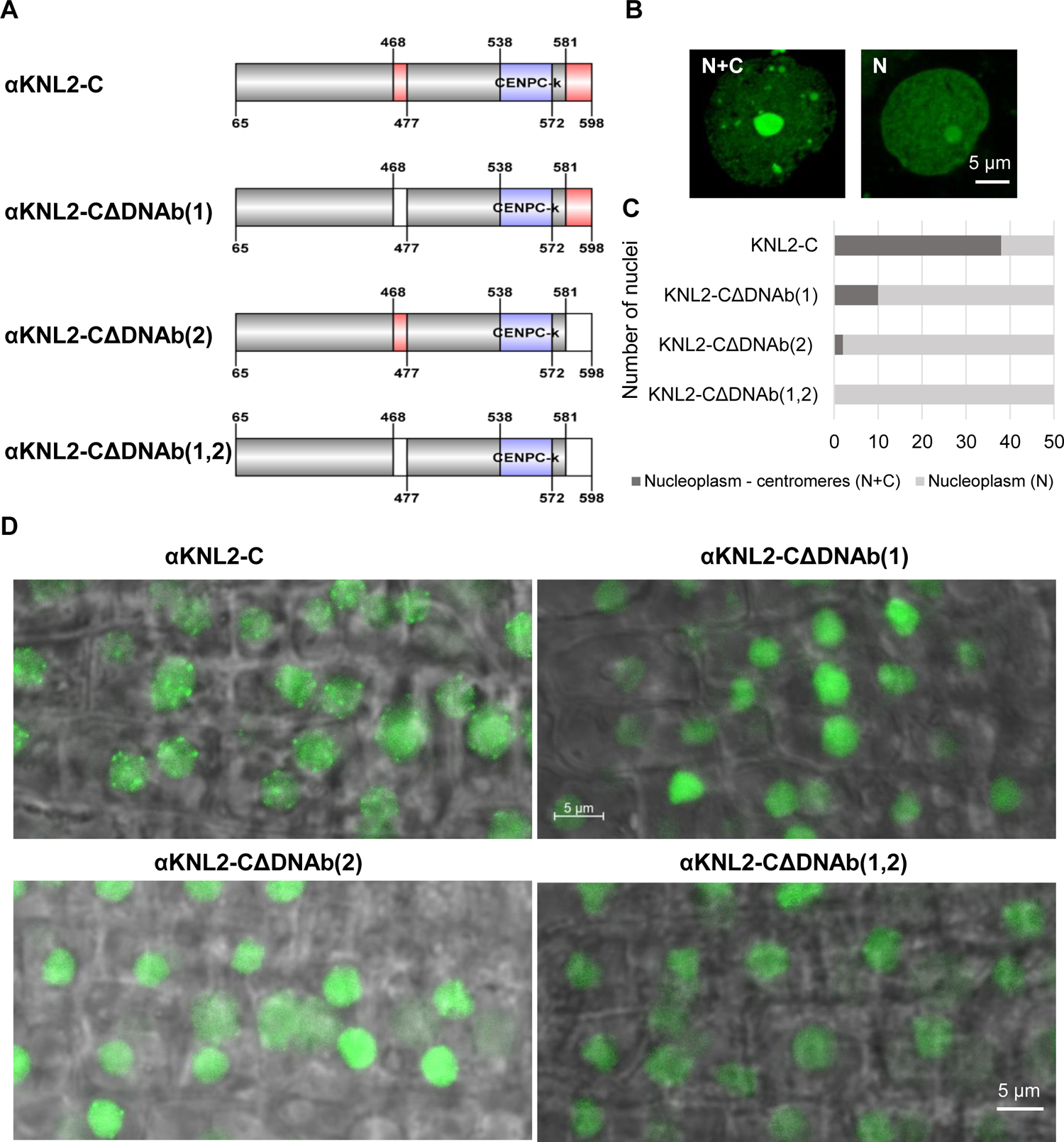
Deletion of DNA-binding regions resulted in reduced centromeric targeting in *N. benthamiana* and *A. thaliana.* **(A)** Schematic representation of the C-terminal part of KNL2 and constructs with deletion of one or two DNA-binding sites. The CENPC-k motif is indicated in purple, DNA-binding sites in red, and deleted sites in white. **(B)** *N. benthamiana* nuclei with fluorescent signals in the nucleoplasm, on presumed centromeres, and in the nucleolus (N+C) and in the nucleoplasm and nucleolus (N). **(C)** the bar chart showing the distribution of nuclei with localization patterns (N+C) and (N) in *N. benthamiana* leaves expressing KNL2 constructs with deletions of DNA-binding regions and an non-mutagenized control. **(D)** Localization patterns of KNL2-C protein fragments with deletions of DNA-binding regions and an non-mutagenized control in root tips of stably transformed *A. thaliana*.

In addition, electrophoretic mobility shift assays (EMSAs) were conducted with αKNL2-C, αKNL2-CΔDNAb (1), αKNL2-CΔDNAb (2) and αKNL2-CΔDNAb (1,2) proteins expressed with TNT SP6 High Yield Wheat Germ and centromeric DNA probe labelled by IR-dye680 (Supplementary Figure 4). Previously, we demonstrated that complete deletion of the CENPC-k motif did not influence the ability of αKNL2 to interact with DNA (Sandmann *et al*., 2017). Initially, we confirmed the interaction between non-mutated αKNL2 (αKNL2-C) and the labelled centromeric *pAL1* probe. Indeed, addition of the unlabelled competitive *pAL1* DNA caused a weaker DNA shift. In contrast to the non-mutated αKNL2-C construct, the αKNL2-C protein fragments with deleted DNA-binding regions showed a reduced *pAL1*-binding ability, with the lowest DNA shift when both DNA-binding motifs were deleted (Figure 6).

**Figure 6.**
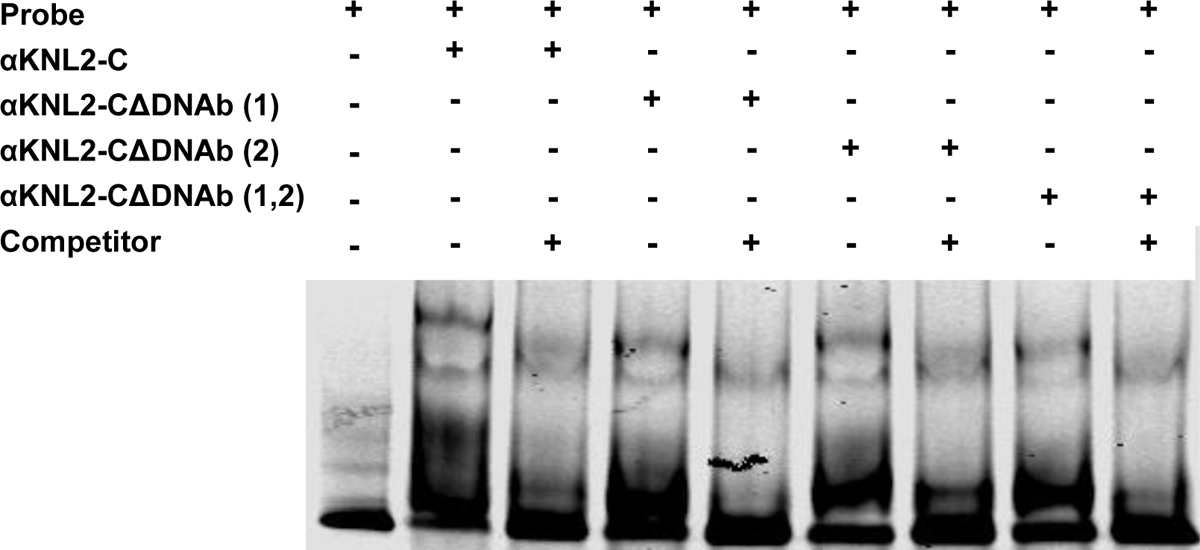
EMSA experiment showing the association of αKNL2-C fragments with the centromeric repeat *pAL1*. Labelled *pAL1* repeat probes were incubated with *in vitro* expressed unmodified αKNL2-C protein or protein fragments lacking one of the DNA-binding sites (αKNL2-CΔDNAb (1)), αKNL2-CΔDNAb (2)) or both sites together (αKNL2-CΔDNAb (1,2)). Competitor experiments with unlabelled *pAL1* indicate specific binding to the *pAL1* DNA probes

### The CENPC-k motif combined with a non-plant DNA-binding region can target centromeres

So far, our findings indicate that the DNA-binding regions of αKNL2 confer DNA binding, while the CENPC-k motif ensures the centromere-specificity of this DNA affinity. Given that DNA-binding regions exhibit lower conservation compared to the CENPC-k motif, but share a positive charge characteristic, we hypothesized that, independent of its sequence, any DNA-binding motif could promote centromere localization, when combined with CENPC-k. In *Escherichia coli*, the abundant histone-like nucleoid structuring protein (H-NS) is known to bind chromosomal DNA at numerous sites, playing a crucial role in bacterial nucleoid organization (Grainger *et al*., 2006). The H-NS protein comprises an N-terminal dimerization domain and a C-terminal non-specific DNA binding domain (Nsdbd) separated by a linker region (Dorman *et al*., 1999). We decided to test whether the Nsdbd domain would lead to anchoring of the CENPC-k motif on centromeres, similar to the endogenous DNA-binding motifs of the αKNL2 protein. Therefore, the Nsdbd was fused to the N-terminus of CENPC-k (Nsdbd-CENPC-k) or flanked on both sides simultaneously (Nsdbd-CENPC-k-Nsdbd) (Figure 7A), and the resulting 3D structures for these fragments were modelled (Supplementary Figure 5A, B). Additionally, a docking analysis of Nsdbd-CENPC-k and Nsdbd-CENPC-k-Nsdbd with centromeric DNA was conducted. Interestingly, the construct containing the CENPC-k motif flanked by two DNA-binding regions exhibited a stronger affinity for centromeric DNA compared to the construct involving a single Nsdbd, with docking and confidence score of −210.31 and 0.76, and −178.53, and 0.63, respectively (Figure 7B, Supplementary Figure 5C). To validate our molecular docking results, Nsdbd-CENPC-k and Nsdbd-CENPC-k-Nsdbd fragments were fused to EYFP at the N- or C-termini, and *Agrobacteria* suspensions expressing the resulting constructs were infiltrated into *N. benthamiana* leaves. The control construct consisting of only the Nsdbd fused to EYFP displayed homogeneous fluorescent signals throughout the nucleoplasm, whereas distinct fluorescent spots at presumed centromeres were additionally observed in many nuclei when one or two Nsdbd were fused to the CENPC-k motif (Figure 7D-E). Immunostaining with anti-NtCENH3 antibodies confirmed the centromeric nature of distinct EYFP signals (Figure 7D).

**Figure 7.**
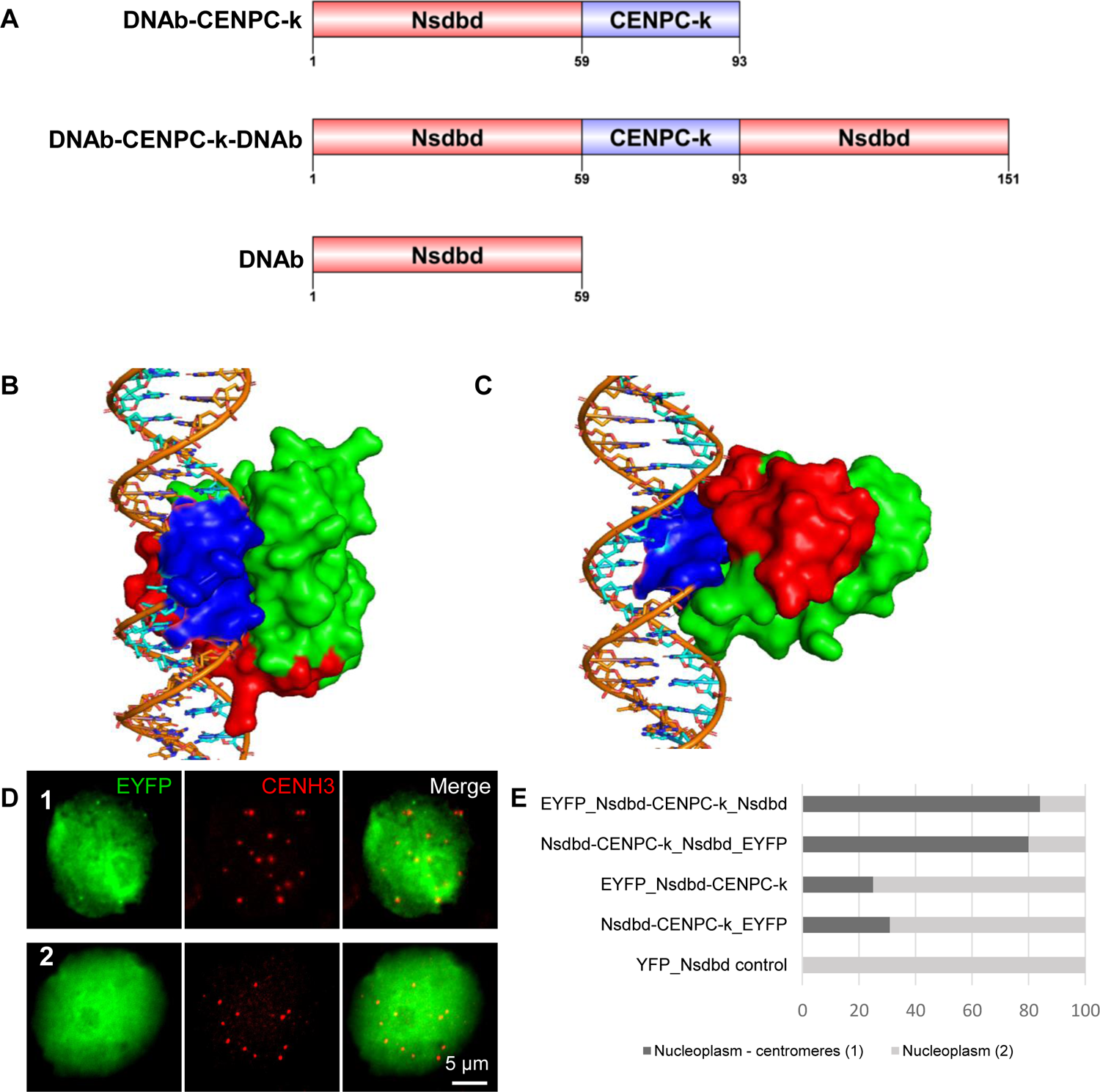
CENPC-k motif in combination with DNA-binding regions of bacteria can target centromeres of *N. benthamiana*. **(A)** Schematic representation of fragments containing the CENPC-k motif and the DNA-binding region(s) of bacteria, as well as the DNA-binding motif alone, used as a negative control. **(B)** The docking of CENPC-k (red) with centromeric DNA (*pAL1*) (orange) showing the interaction via DNA-binding regions of bacteria (green) from both sides. The interaction sites of DNA is shown in surface (blue). **(C)** The interaction sites were reduced between CENPC-k (red) with *pAL1* (orange) when fused with DNA-binding regions on its N-terminal. The protein is shown in surface and the DNA in sticks models. (**D)** Distribution of fluorescence patterns in nuclei of *N. benthamiana* leaves infiltrated with constructs containing CENPC-k in combination with DNA-binding region of bacteria or DNA-binding region alone in fusion with EYFP. The two different patterns of fluorescence signals such as nucleoplasmic and centromeric (1) and nucleoplasmic (2) were defined. Frequency of nuclei displaying these two fluorescence patterns were determined for each construct based on 100 nuclei in *N. benthamiana* leaves. To confirm a centromeric localization of all fusion constructs in *N. benthamiana,* a double immunostaining with anti-GFP and anti-CENH3 antibodies was performed.

## Discussion

The interaction between inner kinetochore proteins and DNA is crucial for proper chromosome segregation during cell division (Cheeseman and Desai, 2008, Musacchio and Desai, 2017). Besides CENH3, CENP-C, CENP-T, CENP-N and αKNL2 are known to interact with DNA (McKinley and Cheeseman, 2014, Sandmann *et al*., 2017). However, the mechanism of this interaction is not well studied. Here we focused on the role of DNA-binding regions of αKNL2 and CENP-C proteins of *A. thaliana* in their centromere targeting mechanism.

KNL2 recruitment to centromeres is a critical step for the kinetochore assembly and in the loading pathway of CENH3. The plant αKNL2 protein contains a conserved SANTA domain at the N-terminus and a centromere-targeting CENPC-k motif at the C-terminus (Lermontova *et al*., 2013, Sandmann *et al*., 2017). Fusion of the C-terminal part of αKNL2 containing the CENPC-k motif, but lacking the SANTA domain, with EYFP showed centromeric localization in transiently transformed *N. benthamiana* and stably transformed *A. thaliana* (Lermontova *et al*., 2013). Since deletion of the CENPC-k motif abolishes centromeric localization of αKNL2-C, it was assumed that this motif plays an important role in targeting of αKNL2 to centromeres. The importance of the CENPC-k motif for centromere targeting has been demonstrated for the KNL2 homologues of chicken (Hori *et al*., 2017) and frog (French *et al*., 2017). Recently using the cryo-EM structure analysis, it was shown that the CENPC-like motif of chicken KNL2 distinguishes between CENH3 and H3 nucleosomes and changes its binding partner at the centromere during the mitotic cell cycle (Jiang *et al*., 2023). The CENPC motif has been identified and described as a typical feature of the CENP-C protein (Brown, 1995, Talbert *et al*., 2004). It is required to direct CENP-C to centromeric nucleosomes in human cells (Song *et al*., 2002, Trazzi *et al*., 2002, Carroll *et al*., 2010). A mutation of the conserved amino acids in the rat CENPC motifs disrupted its ability to bind to centromeric nucleosomes (Kato et al. 2013). Interestingly, the abolished centromeric localization of αKNL2-C, caused by deletion of the CENPC-k motif, can be restored by insertion of the corresponding motif of *Arabidopsis* CENP-C (Sandmann *et al*., 2017).

Here, we showed that the CENPC motifs of αKNL2 and CENP-C alone are insufficient to target centromeres in *N. benthamiana* and *A. thaliana*. Fusion of either CENPC-k or CENPC-m with EYFP resulted in fluorescent signals in nucleoplasm and cytoplasm, but no centromeric localization, neither in transiently transformed *N. benthamiana* nor in stably transformed *A. thaliana*. Previous studies have shown that αKNL2 of *A. thaliana* (Sandmann *et al*., 2017) and the CENP-C protein of maize (Du *et al*., 2010) possess the ability to interact with DNA in a sequence-independent manner. Hence, we hypothesized that the DNA-binding capability might contribute to stabilizing KNL2 and CENP-C at centromeres. Our investigation revealed positively charged regions flanking both sides of the CENPC-k and CENPC motifs. Although no well-defined DNA-binding domains were identified, we speculated that these positively charged regions are involved in sequence-independent DNA-binding of KNL2 and CENP-C proteins. Indeed, using KNL2 or CENP-C protein fragments containing the CENPC motifs and one or both putative DNA-binding regions in fusion with EYFP, exhibited centromeric localization in *N. benthamiana* leaf nuclei. The centromeric localization of corresponding αKNL2 fragments was also observed in *A. thaliana*, but no transformants expressing the CENP-C fragment could be generated. It is possible that overexpression of this region has a toxic effect on the development of transformants, since the resulting protein fragment is able to localise at centromeres and integrate into the corresponding protein complexes, but cannot fulfil the functionality of a full-length protein.

Deletion of one of the DNA-binding regions from the C-terminal part of αKNL2 reduced its centromeric localization, while deletion of both DNA-binding regions abolished it completely despite the presence of the centromere targeting CENPC-k motif. Our experimental data were also confirmed by the bioinformatic prediction of the αKNL2-DNA interaction model. We conclude that targeting of αKNL2 to centromeres requires the CENPC-k motif, but its anchoring at centromeres depends on the DNA-binding regions (Figure 8). Therefore, we postulate that the fusion of the CENPC-k motif with any DNA-binding sites capable of binding DNA in a sequence-independent manner would result in targeting to the centromere. Indeed, fusion of CENPC-k to the non-specific DNA-binding domain (Nsdbd) of a bacterial histone-like nucleoid structuring protein (Grainger *et al*., 2006) promoted its centromeric localization in *N. benthamiana*.

**Figure 8.**
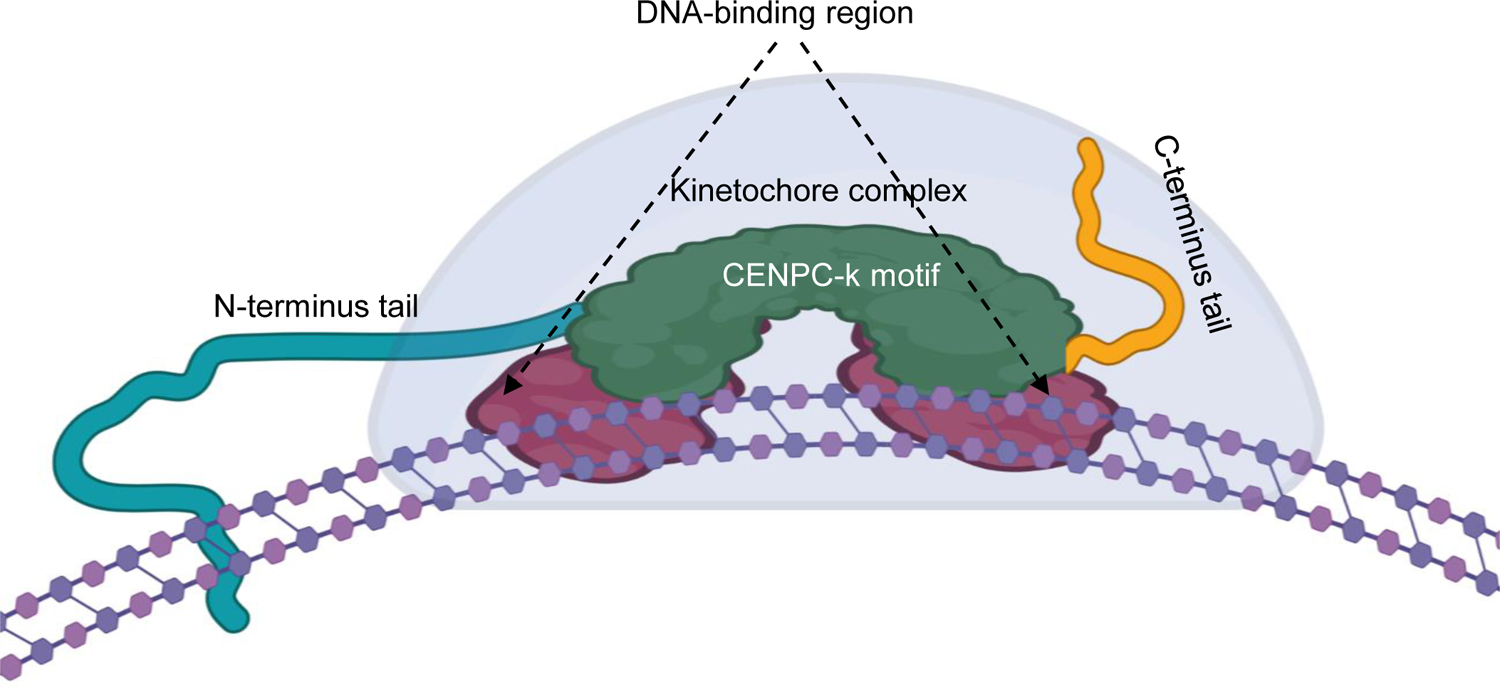
Mechanism of anchoring of kinetochore complex proteins to the centromere through DNA-binding regions.

Furthermore, using online tools, M18BP1/KNL2 sequences from various vertebrates and *C. elegans*, as well as α-, β-, γ-, and δKNL2 variants from plants were analyzed. All proteins tested demonstrated the ability to bind to DNA/RNA nucleic acids as shown in Table 1. Based on this analysis, we suggest that the M18BP1/KNL2 protein from different organisms may use a similar centromere targeting mechanism, where DNA binding sites plays a crucial role in its anchoring to centromeres. A review of Ramakrishnan Chandra *et al*. (2023) described the ability of proteins involved in kinetochore assembly to interact with DNA and RNA in a sequence-independent manner. Apparently, this mechanism allows kinetochore proteins to adapt to the very rapid evolution of centromeric sequences and to form kinetochores in new centromeric regions in the case of inactivation of the original centromere. Sequence analysis of αKNL2 and CENP-C in different plant species revealed that the DNA-binding regions of these proteins are much less conserved than the CENPC motifs. However, a common characteristic of the DNA-binding regions of αKNL2 and CENP-C in all plant species is the enrichment in positively charged amino acids.

## Data availability

All relevant data are within the manuscript and its Supporting information files.

## Acknowledgements

We thank Heike Kuhlmann and Anette Heber for excellent technical assistance and Ingo Schubert for critical reading of the manuscript.

