## Supplementary Figures for "Centromeric localization of KNL2 and CENP-C proteins in plants depends on their centromere-targeting domain and DNA-binding regions"

A

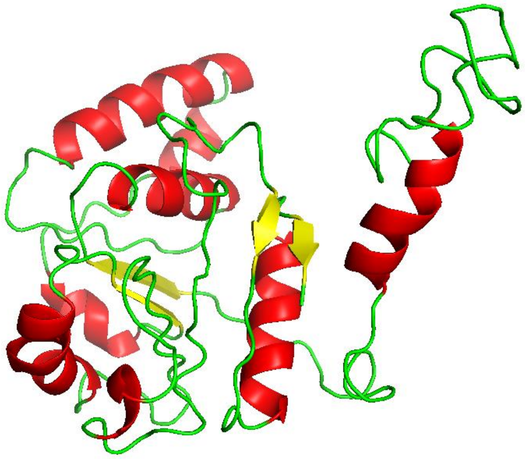

B

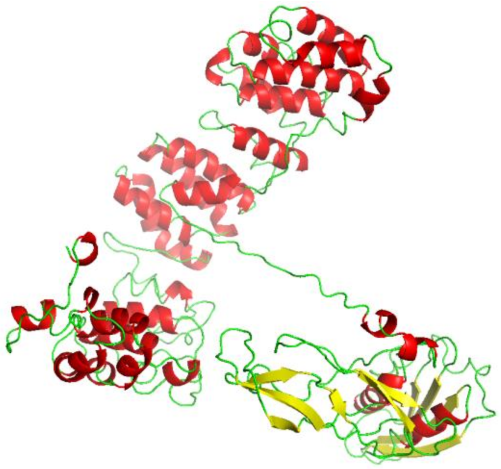

C

| Rank of the predicted model | 1 | 2 | 3 |
|---|---|---|---|
| **αKNL2-C_*pAL1*** | | | |
| Docking Score | -238.2 | -231.9 | -225.9 |
| Confidence Score | 0.85 | 0.83 | 0.82 |
| **αKNL2-CΔDNA-binding (1), (2)_*pAL1*** | | | |
| Docking Score | -179.1 | -165.7 | -163.0 |
| Confidence Score | 0.53 | 0.52 | 0.51 |

D

| Rank of the predicted model | 1 | 2 | 3 |
|---|---|---|---|
| **CENP-C_*pAL1*** | | | |
| Docking Score | -248.4 | -225.2 | -224.4 |
| Confidence Score | 0.87 | 0.81 | 0.81 |
| **CENP-C-ΔCENPC, DNA-binding (1), (2)_*pAL1*** | | | |
| Docking Score | -171.8 | -170.1 | -169.6 |
| Confidence Score | 0.60 | 0.59 | 0.59 |

**Supplementary Figure 1 | The three-dimensional models of αKNL2-C and CENPC proteins and its interaction score with *pAL1***

**(A, B)** The three-dimensional structure of αKNL2-C **(A)** and CENP-C **(B)** protein predicted by I-TASSER. **(C, D)** The table showing the docking and confidence scores of top 3 prediction models for αKNL2-C and αKNL2-CΔDNA-binding (1),(2) **(C)** and CENP-C and CENP-C-ΔCENPC, DNA-binding (1),(2) **(D)** interaction with *pAL1.*

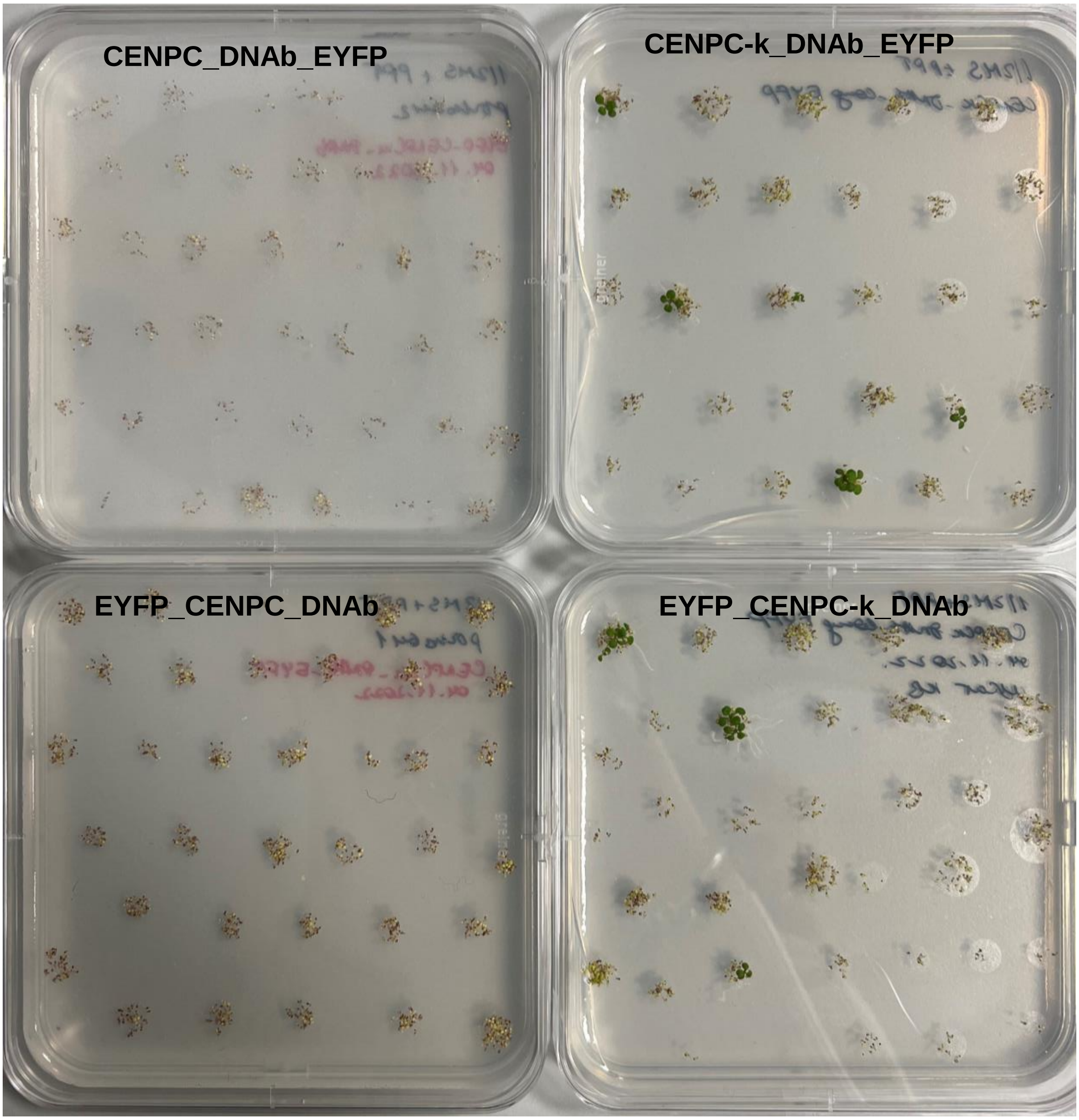


**Supplementary Figure 2 | Germination of seeds of primary transformants (T0) on selective 1/2 MS medium containing 20 µg/ml PPT.**

Despite three repeated transformations of Col wild type plants with CENPC_DNAb constructs (left part), no transformants were selected. In contrast, positive transformants were selected in simultaneously conducted transformations with CENPC-k_DNAb constructs (right part).

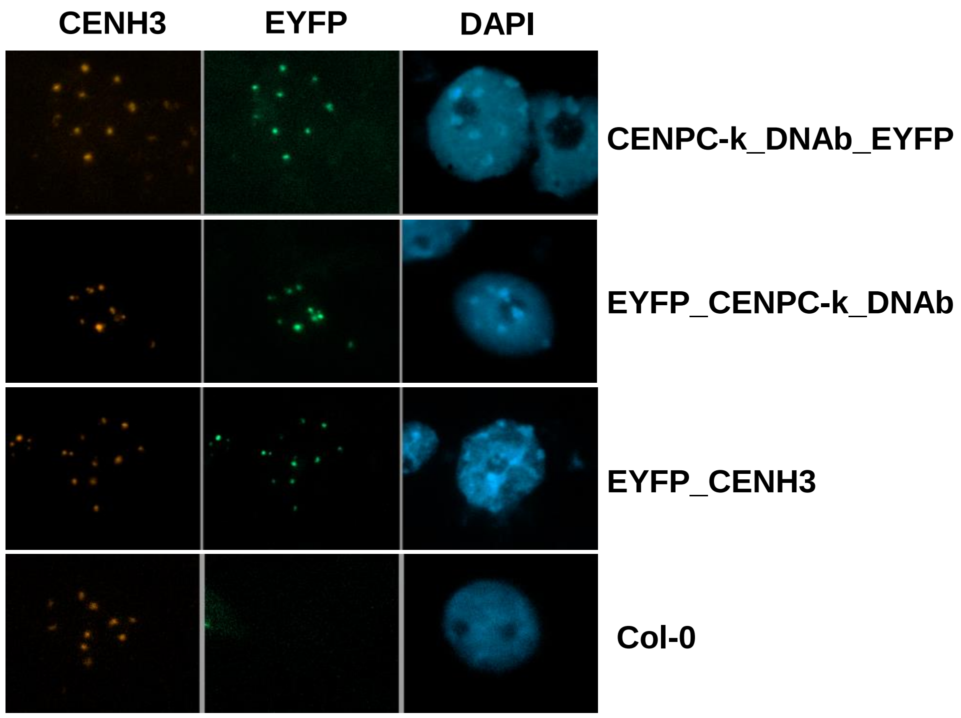


**Supplementary Figure 3 | Double immunolabeling of root meristem nuclei using anti-GFP and CENH3 antibodies**

To verify the centromeric localization of CENPC-k_DNAb fragments fused to EYFP in transgenic *A. thaliana* plants, we conducted double immunostaining with anti-CENH3 and anti-GFP antibodies on root tip nuclei of both CENPC-k_DNAb_EYFP and EYFP_CENPC-k_DNAb transgenic plants. As a positive control for the anti-GFP antibodies, we utilized EYFP_CENH3 transgenic plants (Lermontova et al., 2006), while Col served as the negative control.

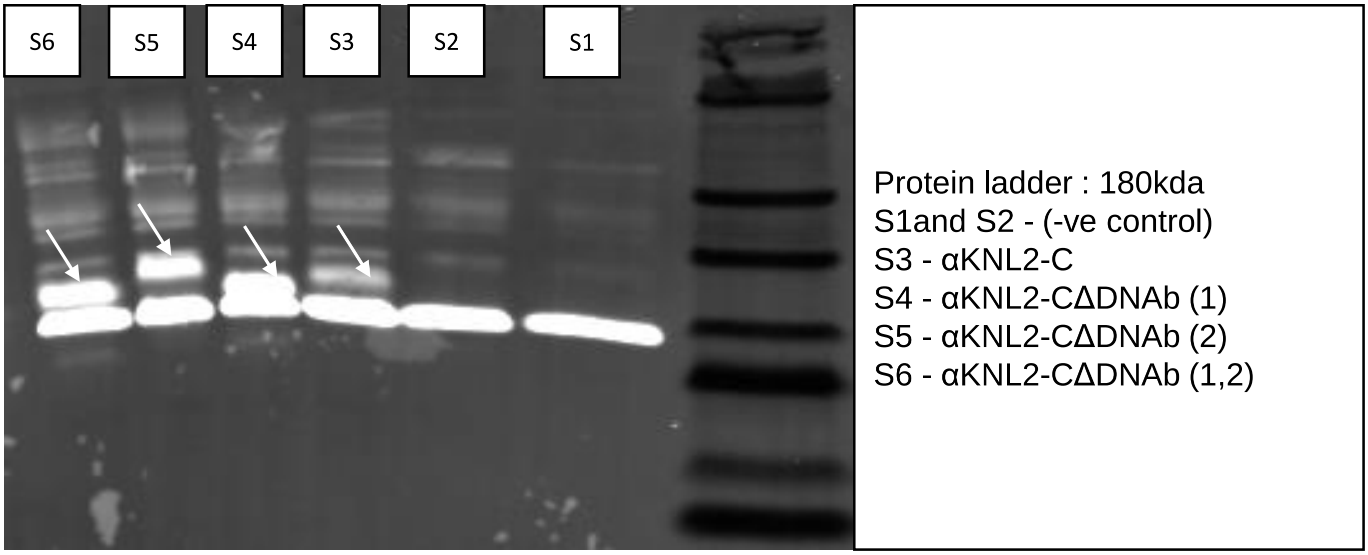


**Supplementary Figure 4 | Western blot analysis to validate the expression of αKNL2-C protein and its mutated variants with deletions of DNA-binding sites in TNT SP6 High Yield Wheat Germ.**

The pF3A WG (Promega) vectors containing the CDS of αKNL2-C, αKNL2-CΔDNAb (1), αKNL2-CΔDNAb (2) and αKNL2-CΔDNAb (1,2) fused with a FLAG tag were used for protein expression with TNT SP6 High Yield Wheat Germ (Promega). Expression of proteins was validated by performing a Western blot analysis against the FLAG-epitope tag using an anti-FLAG antibody. Arrows show αKNL2-C protein variants expressed in TNT SP6 High Yield Wheat Germ system.

A

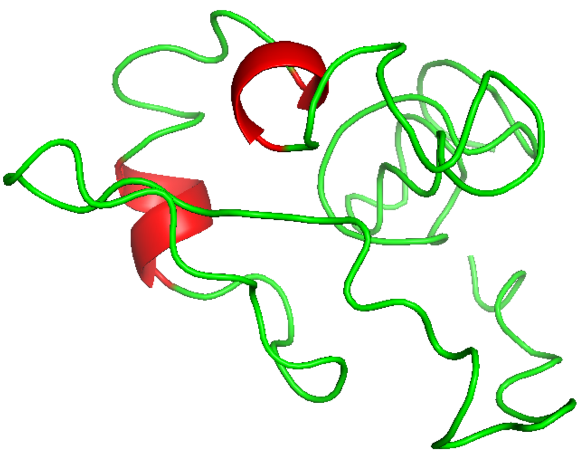

B

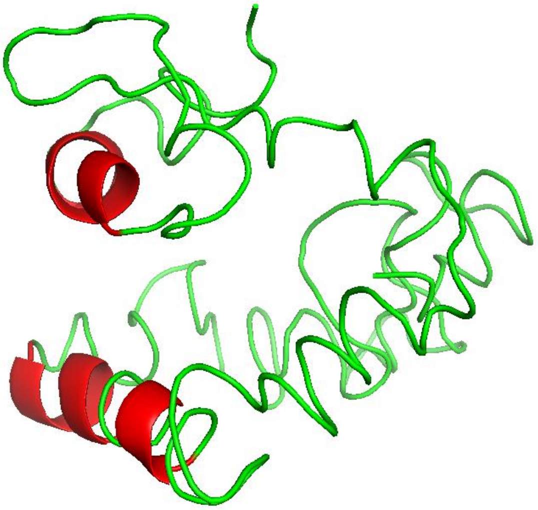

C

| Rank of the predicted model | 1 | 2 | 3 |
|---|---|---|---|
| **Nsdbd-CENPC-k-Nsdbd-*pAL1*** | | | |
| Docking Score | -210.31 | -197.15 | -193.08 |
| Confidence Score | 0.76 | 0.71 | 0.70 |
| **Nsdbd-CENPC-k-*pAL1*** | | | |
| Docking Score | -178.53 | -176.49 | -171.89 |
| Confidence Score | 0.63 | 0.62 | 0.60 |

**Supplementary Figure 5 | The three-dimensional models of non-specific DNA binding domain (Nsdbd) with CENPC-k and its interaction score with *pAL1***

**(A, B)** The three-dimensional structure of Nsdbd-CENPC-k **(A)** and Nsdbd-CENPC-k-Nsdbd **(B)** protein predicted by I-TASSER. **(C)** The table showing the docking and confidence scores of top 3 prediction models for Nsdbd-CENPC-k-Nsdbd and Nsdbd-CENPC-k interaction with *pAL1.*
